# Network-based population analysis of 31,595 gonococcal genomes reveals phase variability, genetic diversity and mobile element dynamics drive antimicrobial resistance and phenotypic diversity

**DOI:** 10.1101/2025.02.28.640638

**Authors:** Duncan Carruthers-Lay, Jessica Lam, Emil Jurga, Scott D. Gray-Owen, John Parkinson

## Abstract

With a highly plastic genome allowing the accumulation of antimicrobial resistance (AMR) determinants, *Neisseria gonorrhoeae* (*Ngo*) is an urgent public health threat. To better understand the spread of AMR in *Ngo* and its ability to persist in humans, we performed a systematic analysis of its global population structure. From 31,595 publicly available genomes, we curated a maximally diverse representative set of 485 gonococcal genomes, which were defined into 12 distinct clades on the basis of shared patterns of single nucleotide polymorphisms. Mapping AMR determinants and mobile elements onto these clades revealed significant correlation with gonococcal population structure. Notably, several variants associated with AMR were identified in genes not previously linked with resistance, suggesting roles as new determinants of resistance or compensatory mutations in resistant strains. Analysis of phase variable motifs, a significant mechanism driving regulation of expression in *Ngo*, identified 57 genes not previously identified as phase variable; *in vivo* and *in vitro* passaging validated our sequence-based definition of phase variability and revealed a strong association of phase variation of the surface proteins Opa and LgtG with improved colonization during infection. Together these findings highlight the ability of systematic comparative genomic analyses to shed new light on the drivers of *Ngo* population structure and identify new AMR determinants.

## INTRODUCTION

The gram-negative bacterial pathogen *Neisseria gonorrhoeae* (*Ngo*) is the causative agent of the sexually transmitted infection gonorrhea. The most common site of gonococcal infection is the urogenital tract, where it typically drives a highly inflammatory response with purulent discharge^1^. Without treatment, infection can result in severe morbidities such as pelvic inflammatory disease or infertility^1^. Since the advent of antimicrobials, *Ngo* has demonstrated the ability to rapidly acquire AMR mechanisms, threatening our ability to treat infections and raising the prospect of untreatable gonorrhea^2^. With rapidly rising rates of infection and widespread development of antimicrobial resistance (AMR), *Ngo* is now considered a high priority pathogen by the World Health Organization, requiring the development of novel therapeutics and additional treatment options^3^.

Facilitating its ability to acquire AMR, *Ngo* is naturally competent with many DNA uptake sequences throughout genome, which enables efficient recognition, uptake and incorporation of genetic matter from gonococcal strains and other pathogenic and commensal *Neisseria* species^4,5^. Driving this “promiscuity” are mobile elements, including plasmids containing AMR encoding genes and the Gonococcal Genetic Island (GGI) which encodes a type IV secretion system, together with several proteins of unknown function^6,7^. Further, many gonococcal genes exhibit phase variability in which replication error-produced changes in the number of tandem repeat sequences can alter protein expression^8^. This genetic promiscuity, phase variability and high rates of recombination support *Ngo’s* ability to rapidly adapt to novel environmental perturbations and result in a highly diverse population structure not observed in the closely related *Neisseria meningitidis*^5^. This population heterogeneity has confounded traditional efforts to define clonal lineages, requiring instead the use of alternative approaches such as multilocus sequence typing (MLST). Complementing such efforts are tools such as PopNet, which take a more agnostic view to selection of loci for determining population structure^9^. Further, PopNet’s use of network graphs help reveal shared genetic elements among multiple strains that are otherwise lost in the rigid structure of more traditional bifurcating phylogenetic trees.

Using genomic data from over 30,000 *Ngo* isolates drawn from all publicly available gonococcal genomes deposited on the NCBI Sequence Read Archive, we sought to apply PopNet to systematically define the global population structure of *Ngo*, investigate its pangenome and assess the incidence of AMR mechanisms and the genetic elements helping support their dissemination. Through this analysis we identify and subsequently validate novel phase variable genes and previously unrecognized AMR associated loci.

## RESULTS

### A representative set of 485 genomes captures temporal and geographic diversity

To begin our analyses of the global population structure of *Ngo*, we first defined a representative set of 485 isolates from an initial set of genome sequence reads from 31,595 gonococcal isolates hosted by the NCBI sequence read archive annotated by year and location of isolation (September 26^th^ 2023; **Supplementary Table S1)**. To maximize geographic diversity, we prioritized isolates based on their continent and year of isolation (see Methods; **Figure 1A**). In brief, following alignment of genome sequence reads to the reference genome (*Ngo* strain TUM19854), representative isolates were selected by grouping isolates by their continent and year of isolation. Isolates within each group were then clustered based on average nucleotide identity estimated by the Jaccard Index, resulting in a set of 451 isolates, to which an additional 20 isolates from rare body sites together with a set of 14 isolates representing the WHO reference collection^10,11^ were added. The resulting reference collection of 485 isolates represent 36 countries, with 38% of the isolates recovered from Canada (72 isolates), the United States (58), and the United Kingdom (54) (**Figure 1B; Supplementary Table S2**). The reference set captures isolates recovered between 1972-2023, with the majority (308, 63.5%) recovered between 2010-2020 (**Figure 1C; Supplementary Table S2**). Notably, our prioritization strategy resulting in a substantially higher proportion (148, 30.4%) of isolates from Asia, South America, and Africa compared to the initial set of 31,595 isolates (**Figure 1D**).

**Figure 1:**
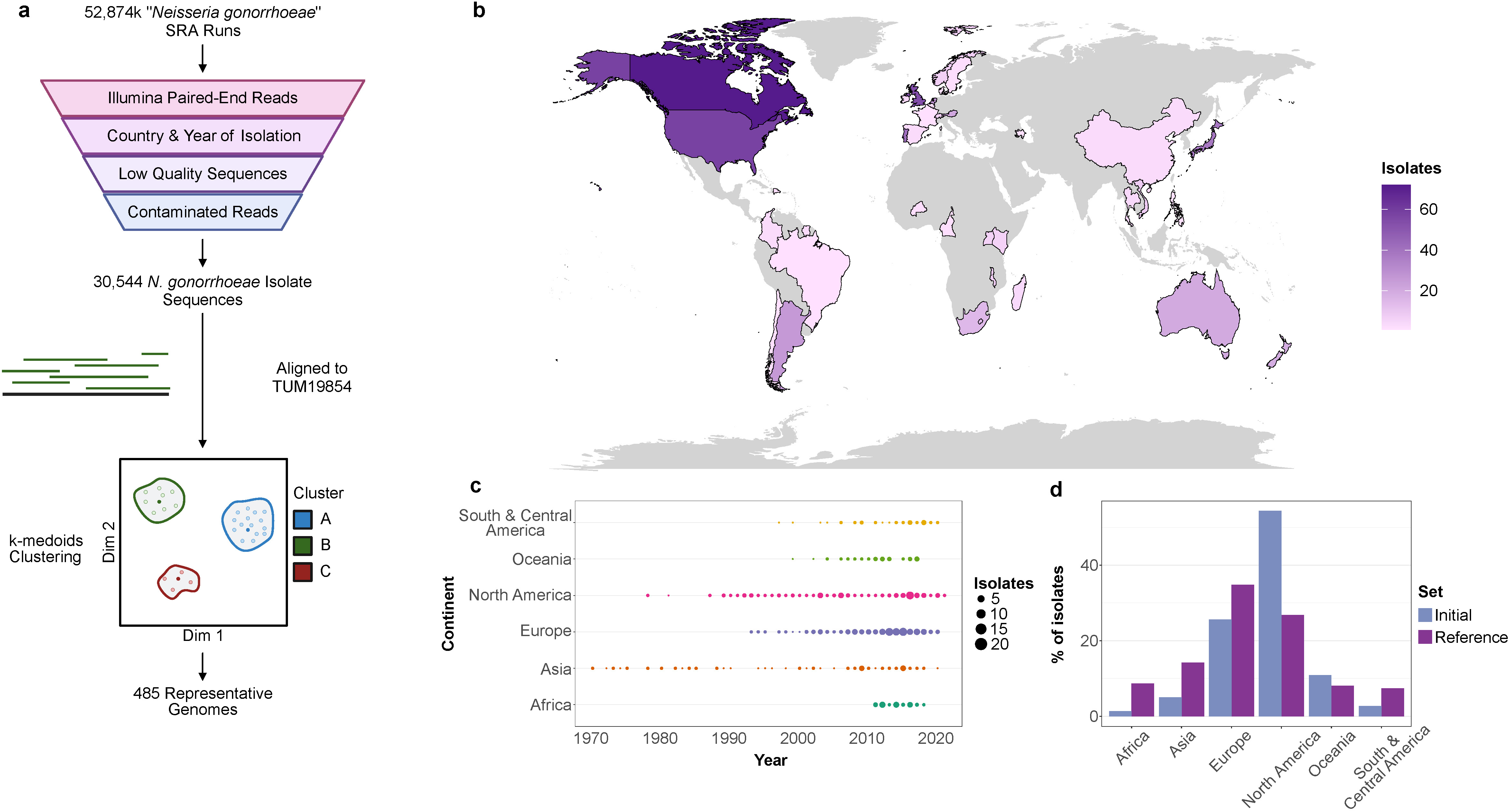
Overview of representative set of 485 *N. gonorrhoeae* genomes. **a** Schematic of selection of representative genomes. **b** Geographic locations of isolates in the representative set. **c** Geographic and temporal distribution of isolates in the representative set. **d** Comparison of the geographic distribution of isolates between the initial set of 52,784 SRA runs and the representative set of 485.

### PopNet defined clades show limited enrichment in specific locales

Having defined our set of representative isolates, we next defined the population structure of these isolates based on shared patterns of single nucleotide polymorphisms (SNPs), relative to the reference genome (*Ngo* strain TUM19854), using PopNet. This revealed 12 clades (designated A-L; **Figure 2A**) ranging in size from 203 isolates (clade A) to 4 isolates (clades K & L). Within these clades we found an enrichment of isolates from Africa (p = 1.77×10^−13^, Fisher’s Exact) and Asia (p=0.00047, Fisher’s Exact) in clade B (**Figure 2Ci**). Clades K and L were exclusively comprised of isolates from Canada, while 4 of 5 isolates from clade J were isolated from Oceania. No other significant associations between continent of isolation and PopNet clades were observed at a significance level of p < 0.05. Together these findings suggest that geography has only a limited impact on *Ngo* diversity, highlighting its ability to rapidly disseminate on a global scale (**Figure 2Bi)**.

**Figure 2:**
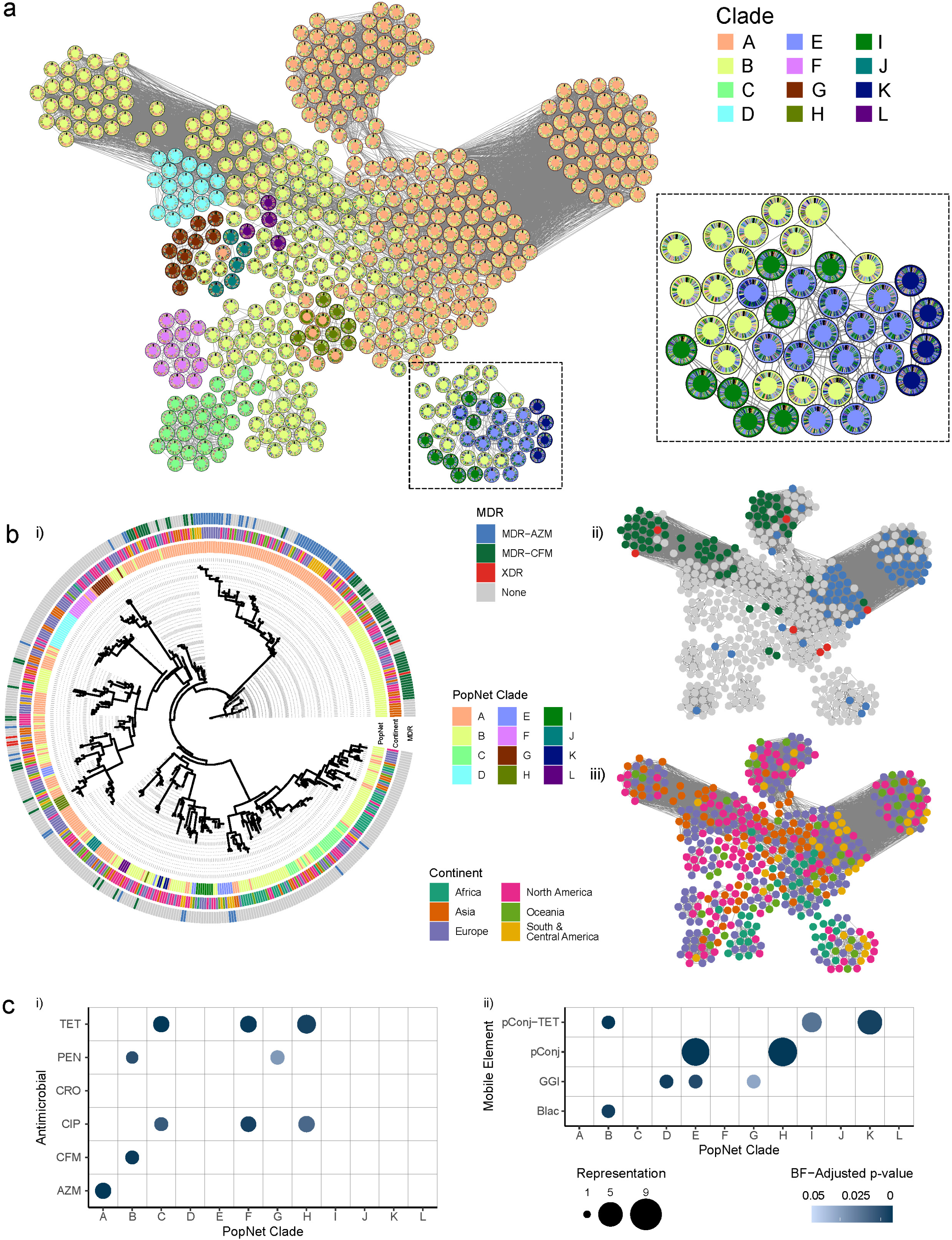
Population structure of *N. gonorrhoeae* is influenced by AMR and the presence of mobile elements. **a** PopNet network view of 485 representative *Ngo* genomes. Nodes represent genomes, coloured by their assigned clades. Annuli show the chromosome painting generated for each genome, with regions sharing a colour showing overlap between genomes. A selection of genomes from clades B, E, I & K are shown to illustrate the chromosome painting visualization. **b** ClonalFrameML generate phylogenetic tree of representative *Ngo* genomes. Outer rings show the assigned PopNet clade, MDR (multi-drug-resistant) status and continent of isolation for each genome. **i**: MDR status overlaid on the PopNet network view. **ii**: Continent of isolation overlaid on the PopNet network view. AZM: azithromycin, CFM: cefixime, XDR: extreme drug resistant. **c** Association between antimicrobial resistance (left, i) or mobile elements (right, ii) and PopNet clades. Only significant associations with a p-value < 0.05 as determined by Fisher’s exact test with Bonferonni multiple testing correction are shown.

Comparison of PopNet clades with a maximum likelihood tree constructed using ClonalFrameML^12^ showed good concordance, with clades broadly remaining intact between the two methods with clear exceptions (**Figure 2Bii**). Comparisons with two other population level clustering methods, cgMLST and PopPunk^13,14^, showed that many cgMLST or PopPunk clusters with more than 5 isolates were located within a larger PopNet cluster (**Supplementary Figure S1**). In the context of a smaller, more diverse set of genomes, the agnostic clustering approach and 2-dimensional network generated by PopNet is better suited to highlight similarities and differences between closely related isolates.

### Pangenome analysis reveals PopNet clades reflect patterns of mobile elements

To compare gene complements across the 485 isolates, we generated genome assemblies *de novo* using unicycler v0.5.0^15^. Genomes were subsequently annotated using Prokka^16^ to identify genes and orthologs defined using Panaroo^17^ (**Supplementary Table S3)**. Across all genomes we identified a total of 2,460 unique genes (i.e. orthogroups), with 1,800 “Core” genes appearing in greater than 95% of isolates, 367 “Shell” genes appearing in between 15% and 95% of isolates and 283 “Cloud” genes appearing in less than 15% of isolates (**Figure 3A**). Accessory (non-core) genes were clustered according to their presence or absence across genomes using the Jaccard index to calculate similarity and MCL clustering with a granularity of 1.10 using Graphia^18^ (**Figure 3C)**. Several large, distinct clusters emerged, which we found to represent mobile elements. These mobile elements were identified through sequence similarity searches against a database of reference sequences generated from the GGI, the *Neisseria* cryptic plasmid, the β-lactamase plasmid, and 3 variations of the conjugative plasmid (pConj; the pEP5050 American tetM plasmid, the pEP5233 Dutch plasmid, and the pEP5289 Dutch tetM plasmid). The GGI was identified in 315 (64.7%) isolates, the cryptic plasmid was identified in 429 (90.1%) isolates, the β-lactamase plasmid was identified in 70 (14.5%) isolates, and the conjugative plasmid was identified in 165 (33.9%) isolates, of which 97 (19.9% of total isolates; 58.8% of pConj containing isolates) were a tetM-containing plasmid.

**Figure 3:**
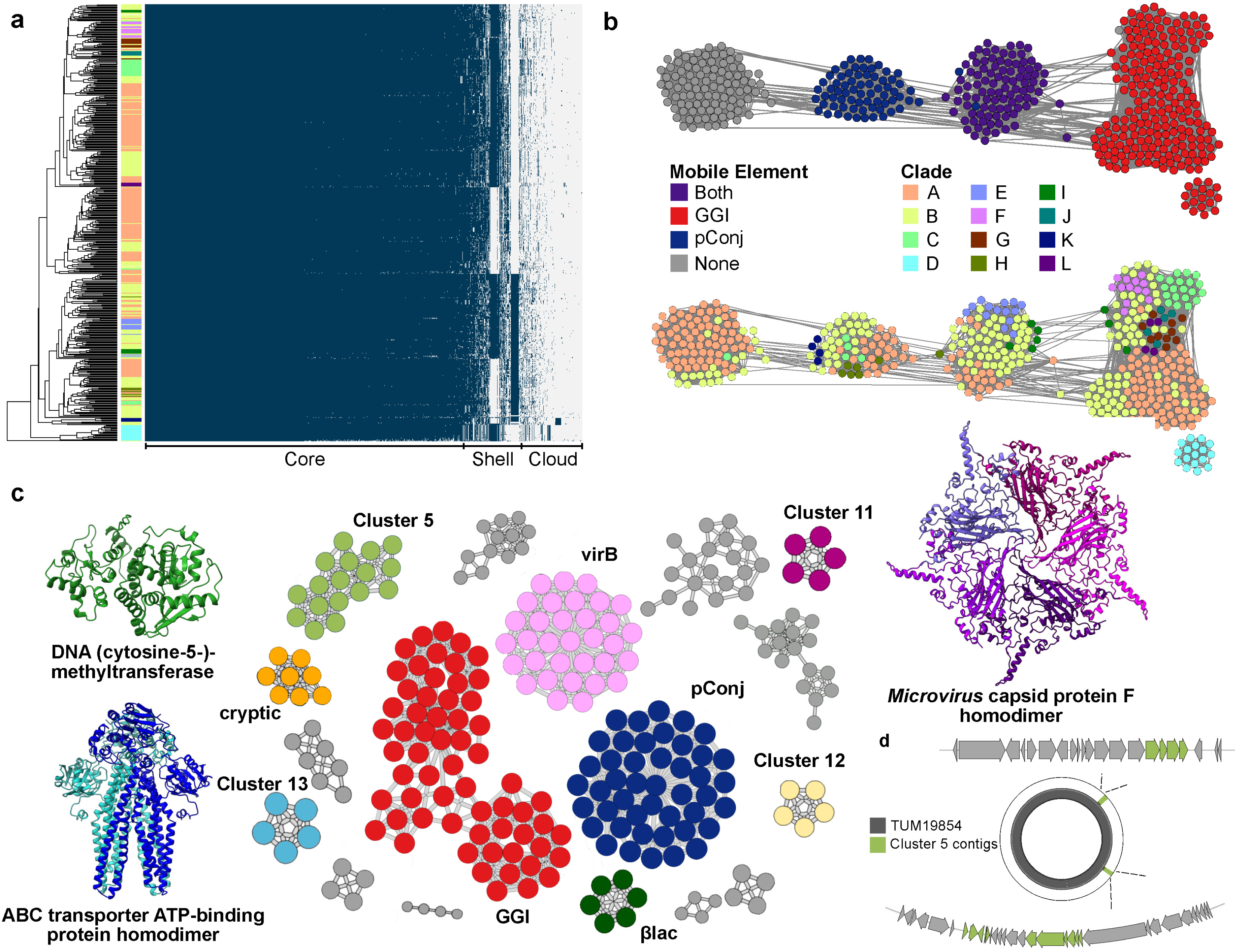
The pangenome of *N. gonorrhoeae* is dominated by large mobile elements, which are associated with PopNet clades determined by chromosomal variants. **a** Pangenome of *Ngo* is composed of 2460 genes, with 1800 Core (present in > 95% of genomes), 367 Shell (present in between 15% and 95% of genomes) & 283 Cloud (present in <15% of genomes) genes. Dendrogram and ordering of genomes on the y-isolates calculated using complete linkage from a Euclidian distance matrix generated based on shared gene content. **b** Top: Network view of isolates clustered by shared gene content coloured by the presence or absence of 2 large mobile elements: the GGI (gonococcal genetic island) and the pConj (conjugative plasmid). Bottom: Genomes coloured by PopNet clade overlaid on the network view of isolates clustered by shared gene content. **c** Clustering of genes based on their co-occurrence in genomes. Clusters defined by MCL clustering as implemented in Graphia which are present in <50% of genomes and composed of 5 or more genes are coloured. Predicted structures from AlphaFold3. **d** Organization of genes from MCL cluster 11 associated with clade D.

Notably, clustering of isolates based on their gene complements was largely congruent with PopNet defined clades (**Figure 3B**). This gene-based clustering appears to have been largely driven by the presence or absence of mobile elements, specifically the GGI and the conjugative plasmid, neither of which are present in *Ngo* TUM19854 and were therefore not included in the PopNet analysis (**Figures 3A and 3B**). Further investigation of PopNet defined clades revealed a spectrum of associations with different mobile elements (**Figure 2Cii**). Of note, isolates from Africa were found to be significantly enriched in pConj-TET (34/42 isolates; p=5.59×10^−19^, Fisher’s exact) and the β-lactamase plasmid (21/42; p=1.34×10^−8^, Fisher’s exact), independent of the year of isolation and likely associated with the continued use of penicillin and tetracycline to treat *Ngo* infections in Africa.

We used the MCL clustering tool implemented in Graphia, to identify clusters of genes based on their shared presence or absence in our set of representative genomes^18^. While some of these clusters lined up with the mobile elements described above, others did not. We sought to further characterize several of these clusters that were present in <50% of our genomes and included at least 5 genes (**Supplementary Table S4**). Of the 5 clusters meeting these criteria, one (MCL cluster 3) represented the previously identified VirB type IV secretion system while the remaining four represent putatively novel mobile elements (**Figure 3C)**. MCL cluster 11 was composed of 5 genes which appear to represent a fully intact *Microviridae* genome present in 7 genomes. All the cluster 11 genes are found on a single contig in all isolates where it was present, and do not appear to be integrated into the gonococcal genome. The largest cluster (MCL cluster 5) was composed of 17 genes present in 17 genomes. Interestingly, all 17 genomes were assigned to PopNet clade D, which includes isolates from the United Kingdom, Japan and the United States, indicating either dissemination within the global gonococcal population or multiple independent acquisitions. Many of the genes present in cluster 5 are annotated as putative phage protein, however these genes do not all appear on the same contig and instead are split between 2 contigs flanked by core genes (**Figure 3D**). The last cluster (MCL cluster 13) was the most common (present in 217/485, 44.7% of our representative set of genomes) and was also composed of 5 genes. Unlike the other clusters, MCL cluster 13 does not appear to be phage related, and appears to include a membrane associated ATP-dependent transporter.

### PopNet defined clades exhibit distinct signatures of AMR mechanisms that are often mutually exclusive

With increased concern over the propagation of AMR within *Ngo* populations, we next used the online platform PathogenWatch^19^ to predict resistance to six antimicrobials – penicillin (PEN), tetracycline (TET), ciprofloxacin (CIP), azithromycin (AZM), cefixime (CFM) and ceftriaxone (CRO) - for each of the 485 reference isolates (**Supplementary Figure S2**). Incidence of resistance to the three older antimicrobials (PEN, TET & CIP) were higher than rates of resistance to the other three and were significantly enriched in isolates derived from Africa (p=0.029, 0.00011 and 0.016 for PEN, TET & CIP respectively, Fisher’s exact).

We found several PopNet clades were significantly associated with resistance to these specific antimicrobials (**Figure 2B).** Beyond the presence/absence of resistance to certain antimicrobials, we also observed clades associated with specific mechanisms of resistance. Clade C was enriched in isolates resistant to TET (23/25 isolates; p=1.63×10^−5^, Fisher’s exact) despite the absence of the pConj-TET plasmid. Instead, resistance appears a consequence of a double mutation (*mtrR* promoter −5delA deletion & *rpsJ* V57M). This is contrasted by the high rates of TET resistance in clade B which also carried the pConj-TET plasmid at rates. The association of AMR patterns and mechanisms of resistance with clades suggests that AMR helps shape the global population structure of *Ngo*.

### Analysis of repeat expansion distributions identifies 57 potentially novel phase variable genes

Previous studies have shown that many *Ngo* genes contain repeats that can result in slipped strand mispairing or site-specific recombination altering their expression and providing a reversible process to adapt to rapidly changing environments. Within our collection of 485 reference isolates we searched for candidate phase variable (PV) genes using the repeat motif finding algorithm PERF^20^ to detect simple sequence repeats. From this set we identified 85 genes with repeats that varied in length, with each repeat length appearing in at least 1% of available sequences for that gene (**Supplementary Table S5**). These genes were compared with a previous set of 34 genes, identified through comparisons of locus tags between the reference strains FA1090 and NCCP11945, with 28 appearing in both sets. This overlap included 11 *opa* genes, 6 genes characterized as *pilC2* or pilin cassettes, and other well-characterized phase variable genes such as *lgtG*, *fetA* and *modA/B*, helping to validate this approach (**Supplementary Table S5**).

The remaining 57 genes represent putative phase variable genes, with the 10 genes exhibiting the greatest repeat length variation shown in **Table 1**. Among these 10, group_1317 is annotated as a cytosine-specific methyltransferase, consistent with other gonococcal methyltransferases (e.g. *modA* and *modB*) known to be phase variable. Another 2 predicted genes (*argS* and *glyS*) are annotated as arginyl- and glycyl-tRNA synthetases, a system which was previously identified in a computational screen for phase variable genes^21^. We identified 2 genes (group_1584 and *prn*) that were annotated as pertactin or autotransporters, with *prn* appearing to encode a near full length autotransporter while group_1584 represents a truncated form, with only the N-terminal domain encoded^22^. Another autotransporter, *autA* is known to be phase variable, however the identified repeat for *autA* is AAGC, while we identified poly-A tracts in group_1584 and *prn*^23^. While all the genes so far described are chromosomally encoded “Core” genes, we also identified a gene (*kfrB*) carried on the conjugative plasmid that possesses a poly-G repeat tract immediately after the start codon; the protein this encodes plays a role in plasmid partitioning^24^.

**Table 1:** Predicted new phase variable genes.

| Gene | Repeat | No. of repeats | Annotation | NGK locus | NGO locus |
| --- | --- | --- | --- | --- | --- |
| group_1584 | A | 33 | Pertactin | NGK_2016 | NGO_1158 |
| group_280 | AACGC | 24 | Hypothetical protein | - | NGO_0366 |
| *argS* | AAGC | 18 | Arginyl-tRNA synthetase | NGK_0832 | NGO_0963 |
| group_1317 | AACGC | 17 | C-5 cytosine-specific DNA methylase | NGK_0522 | NGO_0365 |
| group_1449 | AT | 11 | Spermidine synthase | NGK_2156 | NGO_1752 |
| group_1206 | C | 11 | ISXO2-like transposase domain | NGK_1147 | NGO_1157 |
| *kfrB* | C | 10 | - | - | - |
| *glyS* | C | 9 | Glycyl-tRNA synthetase beta | NGK_2629 | NGO_2154 |
| *prn* | A | 8 | Autotransporter | NGK_1206 | NGO_0694 |
| *yigL* | C | 7 | Hydrolase | NGK_1967 | - |

### Passaging gonococci *in vivo* reveals phase variable gene selection

We aimed to evaluate the potential impact that phase variable genes would have on enabling gonococci to adapt to and survival in a more natural environment. To accomplish this, we utilized a lower genital tract colonization model with female FVB mice expressing human CEACAM3, CEACAM5, and CEACAM6^25^. A schematic of the mouse passaging work is shown in Figure 4A.

**Figure 4:**
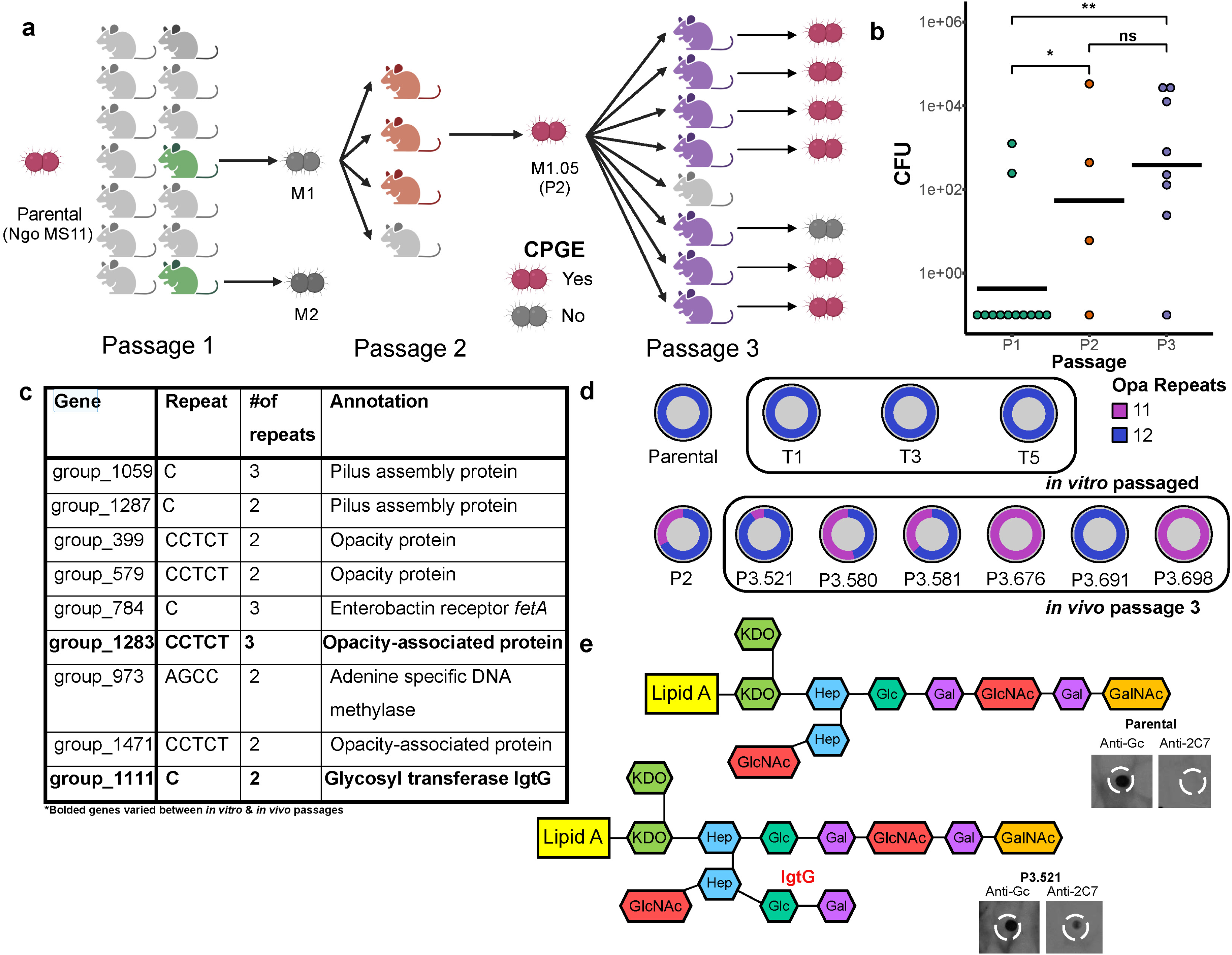
Passaging of *N. gonorrhoeae* improves colonization in a mouse model of infection which may be associated with phase variation of *opa*. **a** Schematic overview of *in vivo* and *in vitro* passaging experiment. Coloured mice represent mice successfully colonized by *Ngo* 2 days post infection. Coloured gonococci indicate isolates sequenced by capillary electrophoresis to confirm the repeat lengths of *opa*. **b** Recovered CFU (colony-forming units) of *Ngo* recovered 2 days post infection. Black bars represent the mean CFUs recovered for each passage. **c** Predicted phase variable genes from mouse passaging experiment. **d** Ratio of *opa* repeats in each isolate sequenced by capillary electrophoresis. **e** Schematic of *Ngo* lipooligosaccharide with (bottom) and without (top) lgtG expression, resulting in the production of the 2C7 epitope. Isolates passaged *in vivo* express lgtG, while no expression is observed in isolates passaged *in vitro.* Immunoblot assay for the 2C7 isotope confirms the expression of lgtG in a passage 3 isolate, which is absent from the parental isolate. KDO: 3-Deoxy-D-manno-oct-2-ulosonic. Hep: Heptose. Glc: Glucose. Gal: Galactose. GlcNAc: N-acetylglucosamine. GalNAc: N-Acetylgalactosamine.

For initial infection (Passage 1; P1), 12 mice were each inoculated with 10^7^ bacteria of N2009 (laboratory-passaged MS11 derivative; denoted parental strain), and vaginal washes were performed every two days to recover viable bacteria. On Day 2, only 2 of the 12 mice remained colonized by the bacteria; these were denoted as M1 and M2 (Figure 4A). Mice infected from the M1 lineage demonstrated progressively higher rates of colonization, 75% and 88% in Passage 2 (P2) and Passage 3 (P3), respectively (**Supplementary Figure S3**). Moreover, we also observed a significant increase in bacterial burden recovered from the P3 isolates compared to the initial inoculum (Wilcoxon ranked-sum, p= 0.0047) (**Figure 4B**). These findings suggest that gonococcal isolates passaged *in vivo* are better adapted for survival during the initial stages of infection **(Figure 4B**).

To differentiate mutations that confer a fitness advantage *in vivo* from those selected by simple subculture, we performed complimentary *in vitro* passaging using the parental strain. From this complementary screen, we identified 9 genes with signatures of phase variability including the known phase variable genes *pilC2*, pilin, *opa* (4), *fetA,*, *lgtG* and *modA* (**Figure 4C**). We observed that the predicted number of repeats in group_1283, which encodes an Opa protein, dropped from 12 to 11 in P2, P3_580 and P3_698. No shift was observed in any of the *in vitro* strains, suggesting that this shift might contribute to the improved colonization of *in vivo* passage 3 isolates. Due to the inherent population heterogeneity in *Ngo*, especially when it comes to phase variable genes, we used capillary gel electrophoresis to determine the bacterial population proportion containing 11 or 12 repeats based on peak density. We found that the parental strain and the 3 tested *in vitro* passage strains showed no variation in repeat length, remaining at 12 repeats the entire time, while the passage 2 and 3 strains were more heterogeneous, with approximately 32.5% of the recovered passage 2 isolates and 100% of P3_676 and P3_698 at 11 repeats (**Figure 4D**). We were unable to obtain sufficient genetic material from recovered P3_680 for analysis. These findings indicate that the majority of the *in vivo*-passaged population has been selected to express this Opa variant (with 11 repeats), consistent with its key role in adhesion to host CEACAMs, though a subset of the population remains Opa off in the tissues^25^.

The selection within *lgtG* (group_1111) was also notable, as the parental and all the *in vitro* strains contained a 10bp-long poly-C repeat, while the P2 and all the P3 isolates shifted to an 11bp long repeat. This shift enables complete transcription of *lgtG*, resulting in the attachment of the β-chain to the gonococcal lipooligosaccharide and production of the 2C7 epitope (**Figure 4E**)^26^. Due to the challenges of differentiating single nucleotide length repeats with capillary electrophoresis, we did not perform this analysis for *lgtG*, however we did confirm the presence of the 2C7 epitope exclusively in the *in vivo* passaged strains through an immunoblot assay. Together these findings highlight the ability of our method to identify phase variable genes in an *in vivo* setting, and also identified phase variation in 2 genes, *opa* and *lgtG*, which may play a substantial role in improving *Ngo*’s ability to colonize and infect a humanized mouse model.

### Analysis of sequence variants reveals that novel sites of genes involved in AMR are under positive selection

In the previous sections our genomic surveys identified several phase variable genes that may help *Ngo* adapt to environmental perturbations. Another signature of genes supporting adaptation is whether they are under positive selection. To identify such genes we used the BUSTED algorithm as implemented in HyPhy^27,28^. Of the 1,853 core genes, 1,724 were sufficiently variable to test for selection. We identified 355 genes with significant evidence for positive selection (likelihood ratio test statistic > 0, p-value < 0.05). As expected, given their direct contact with the environment, positively selected genes were significantly enriched in outer membrane proteins (Benjamini-Hochberg (BH) corrected p-value 0.00615) (**Figure 5A**). In addition, we also identified several AMR-associated genes as being under positive selection, including *porB*, *penA*, *parC*, *mtrD*, and *gyrA*. To identify specific sites in AMR genes under positive selection, we applied the FUBAR algorithm as implemented in HyPhy^29^. Sites with a posterior probability of positive selection > 0.9 were considered as sites with strong evidence for selection.

**Figure 5:**
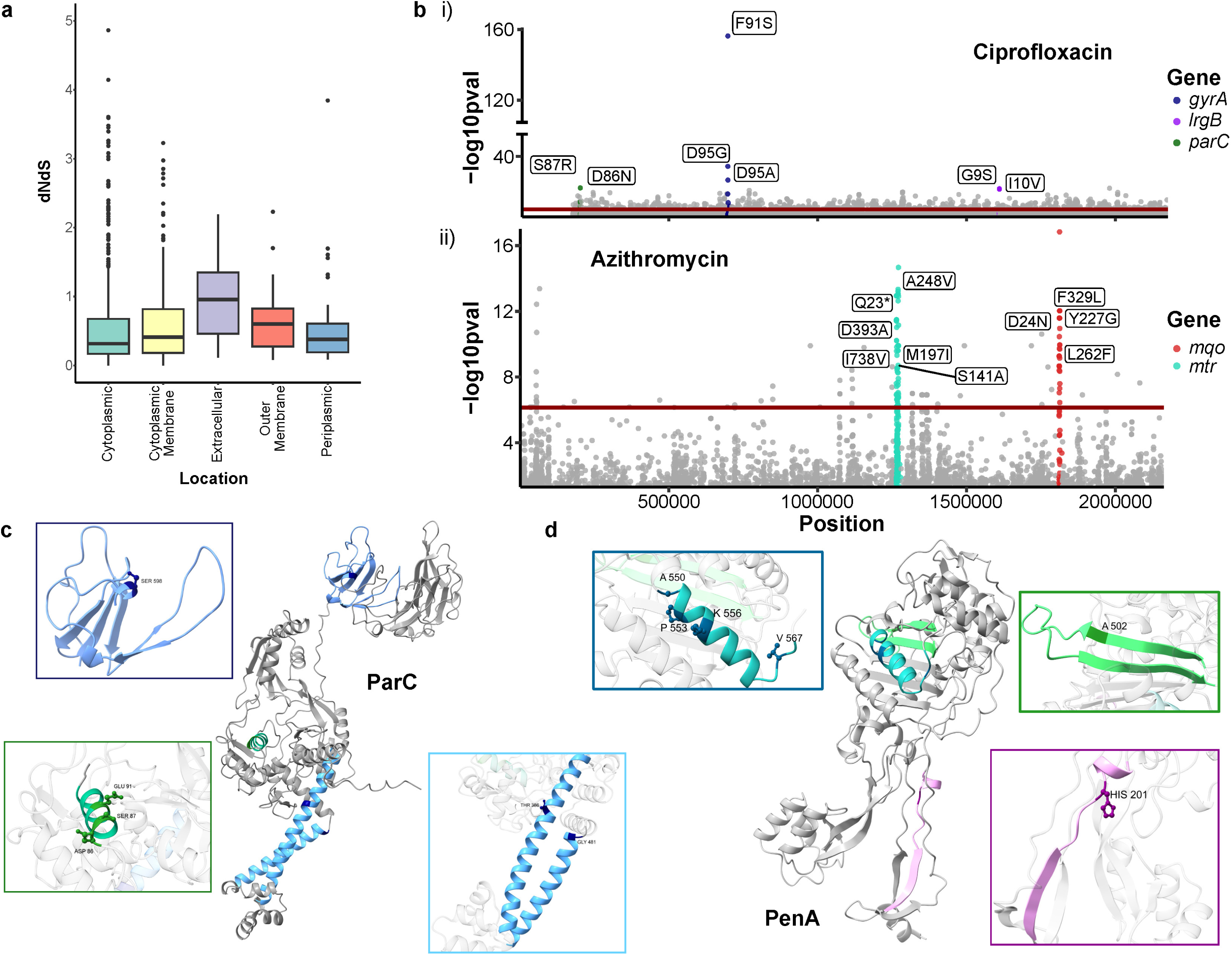
GWAS and tests for positive selection identify sites linked to AMR in *N. gonorrhoeae*. **a** Boxplot of dN/dS ratio for positive selection of *Ngo* genes by predicted cellular localization. **b** Manhattan plots of variants with significant associations with ciprofloxacin (top) and azithromycin (bottom) resistance. Only variants with an LRT p-value <0.05 are shown. Red lines indicate significance thresholds determined by Bonferroni correction with the number of unique variant patterns as the number of multiple tests. **c** Predicted residues with strong signatures of positive selection in *parC*. Residues highlighted in green are under positive selection and known to contribute to ciprofloxacin resistance. Residues in blue are under positive selection with no known links to AMR. Structural prediction from AlphaFold3 visualized with USCF ChimeraX. **d** Predicted residues with strong signatures of positive selection in *penA*. Highlighted residues are under positive selection with no known links to AMR. Structural prediction from Alphafold3 visualized with USCF ChimeraX.

Focusing on the gyrase subunit GyrA, our analyses identified residues S91 and D95, previously characterized as conferring resistance to ciprofloxacin^30^, as being under positive selection (**Supplementary Table S6**). Ciprofloxacin resistance is also associated with mutations in residues D86, S87, S88, and E91 in the type IV topoisomerase subunit, ParC. In addition to sites D86, S87 and E91, which interact directly with ciprofloxacin^31,32^, we also identified residues T386, G481, and S598 as being under positive selection. Residues T386 and G481 reside in the α-helices that connect the catalytic and dimerization domains, while residue S598 is located in the C-terminal domain (**Figure 5B**)^33^. Independent of these, we identified 8 residues (310, 642, 742, 922, 926, 1052, 1065, 1067) with signatures indicating strong positive selection in *mtrD*, likely reflecting the acquisition of mosaic alleles from other *Neisseria* sp. associated with azithromycin resistance^34^. For the known penicillin resistance gene, *penA*, we identified 6 residues with strong signatures of positive selection (**Figure 5C**). We identified residues A550, P553, K556, and V567 within the α11 helix, which also includes the P551 variant previously linked to penicillin resistance through inhibition of acylation by β-lactamases^35^. We also identified a residue (A502) in the region between the β3 and β4 loops that has been linked to ceftriaxone resistance^36^. The final residue under positive selection was Y201, which is located between the α3n helix and β6n loop in the N-terminal domain of PenA, which to date has not been linked to AMR^35^. Cumulatively, these finding reveal signatures of strong positive selection in individual residues of AMR-associated genes, which may represent new or complementary sites contributing to resistance. This reflects prior work that has used signatures of positive selection to identify genes contributing to antimicrobial resistance in *Escherichia coli*^37^.

### Genome wide association study identifies previously unreported sequence variants associated with AMR

Beyond signatures of positive selection, genome wide association studies offer a powerful approach to identify SNPs associated with specific traits such as AMR^38^. Here we used the software tool, pyseer, with the linear mixed model mode to identify SNPs significantly associated with predicted resistance to antimicrobials (**Supplementary Table S7)**^39^. We identified several well-known variants together with several less characterized mutations significantly associated with AMR. For ciprofloxacin resistance, the former includes the well studied S91F and D95A/N mutations in GyrA, as well as an S87R mutation in ParC (**Figure 5Di).** However, we also identified two mutations (G9S & I10V) in LrgB, a cytoplasmic membrane-associated murein hydrolase transporter, as being associated with ciprofloxacin resistance. We found multiple known and uncharacterized variants with *penA* associated with resistance to penicillin, ceftriaxone and cefixime (**Supplementary Figure 4A, 4Ci & 4Cii).** For tetracycline resistance, our analysis identified a variant in the known resistance gene, RpsJ (M57V), as well as multiple mutations in *rsmB* which encodes a methylase that interacts with the target of tetracycline, the 16S portion of the small ribosomal subunit (**Supplementary Figure 4B**). Finally, we identified multiple significantly associated variants associated with azithromycin resistance in genes associated with the MTR multi-drug efflux pump (*mtrC, mtrD,* & *mtrE*) (**Figure 5Dii**). Interestingly, we also identified multiple variants in *mqo*, a malate-quinone oxidoreductase involved in the TCA cycle to be significantly associated with azithromycin resistance. Azithromycin is not known to target *mqo* or the TCA cycle, and pairwise structural alignments from AlphaFold predicted structures using TM-align show *mqo* shares minimal similarity with *rplV* (TM-score 0.11, 7% identity) or *rplD* (TM-score 0.17, 6% identity), known ribosomal targets of azithromycin, indicating if a direct interaction exists it must be unique^40–43^ (**Figure 5E)**. Together these findings reveal the potential of genetic analyses to reveal novel AMR determinants within and beyond known AMR-associated genes.

## DISCUSSION

*Neisseria gonorrhoeae* are not generally considered to be a clonal species based upon classical phylogenetic analyses^5^. Here, we have used PopNet to analyse the population structure based upon localized patterns of SNPs to consider whether temporal, geographic, or clinical metadata related to diversity. We also sought to determine whether AMR or other genetic determinants have a previously unrecognized distribution. Our set of 485 representative genomes successfully captured the genetic diversity of the larger set of 31,595 isolates, increasing the relative contribution of isolates collected from outside North America and Europe. Ensuring representation across the globe is critical if we are to understand the impact of global public health programs regarding surveillance and treatment options, factors that are acutely elevated for a pathogen that is restricted to a single host^2^. Applying the population genomics tool PopNet, we define 12 clades, which, with the exceptions of clades B, K and L, contain isolates that are globally dispersed, suggesting other, non-geographical, factors drive gonococcal population structure. While different, we did observe a high degree of overlap between clades defined by other tools (cgMLST & PopPunk)^13,14^, supporting the utility of PopNet to define genetically distinct sub-populations.

Given the lack of overlap between the geographical location of isolation and clade membership, we investigated how global population structure may be influenced by other factors. Most identified clades were not significantly associated with other available metadata, including sex of patient, site of isolation, year of isolation and country or continent of isolation. We then investigated other factors not captured in isolate metadata: AMR and the presence of mobile genetic elements. In keeping with previous findings, we observed PopNet-defined clades correlate with a strong connection between the presence of specific mobile elements and AMR determinants^44,45^. The association with mobile elements is particularly striking as PopNet clustering only accounts for genetic variation in core chromosomal genes, revealing an association between lineages and specific mobile elements. From these findings, it appears that AMR and mobile elements are the primary drivers of the gonococcal population structure as opposed to traditional metadata like geographic location or site of isolation. Clade B appears to be an exception, as it is significantly associated with isolates recovered from Africa and Asia. Hoewver, this association can be explained by the elevated presence of the β-lactamase plasmid and the TET-containing conjugative plasmid in group B. This is in line with previous studies that have suggested that isolates from lower and middle income countries are more likely to carry plasmids encoding resistance to penicillin or tetracycline^46^.

Phase variability is a key mechanism used by *Ngo* to evade the immune system by independently switching on or off (or up-/down-regulating) protein expression of a variety of different genes^22^. While extensive work has been done to characterize these genes in reference strains, and/or by identifying homologs in the closely related species, *N. meningitidis*, recent evidence has suggested that the phase variable repertoire of *Ngo* could be much larger than previously thought^21,22^. In our analysis, we identified 57 genes containing tandem repeats of varying lengths which represent putative phase variable genes. To complement our population-based approaches, we performed an *in vitro* and *in vivo* passaging experiment in an attempt to drive the emergence of phase variability in the reference gonococcal strain MS11. Passaging of gonococci *in vitro* has previously been used to identify phase variable genes^10,27^, however this strategy likely selects for expression of factors required for tissue colonization to phase “off”. Our *in vivo* passaging in the lower genital tract of CEACAM-humanized mice aimed to identify phase variable genes that help support *Ngo*’s ability to infect and propagate within this mucosal niche. During the passaging experiment, we observed improved colonization of gonococcal isolates with each successive passage, indicating adaptation to the mouse host. Through this tandem approach, we observed several previously characterized phase variable genes (*pilC*, *opa, modA, lgtG* & *fetA*) to phase vary during passaging, of which two (group_1283, *opa* & group_1111, *lgtG*) varied in repeat number only in the *in vivo* passaged isolates. We confirmed the change in repeat number within the Opa protein encoded by group_1283 of *in vivo* passaged 2 and 3 isolates; since this turns on the expression of this adhesin, it likely contributes to the observed improvement in colonization^47^. We also observed a shift in the number of repeats in all *in vivo* passage 2 and 3 isolates, resulting in transcription of *lgtG*, with no shift observed in the *in vitro* passaged strains. Expression of LgtG leads to the production of the 2C7 epitope, which is present in a high proportion of clinical isolates, indicative of a role in gonococcal pathogenesis and colonization^48^. The selection for LgtG expression exclusively in all *in vivo*-passaged isolates supports this conclusion, and demonstrates the utility of passaging and sequencing for identifying genomic variation linked to improved gonococcal pathogenicity.

Given the genomic plasticity of *Ngo* and selection constraints imposed by the host immune system, we hypothesized many *Ngo* genes may be subject to positive, diversifying selective pressure. We identified 355 of 1,853 core genes to be under positive selection. This number dramatically expands on the previous estimate of only 10 *Ngo* genes under positive selection, likely reflecting differences in methodology as well as the larger number of genetically diverse isolates used here^49^. We found no functional enrichment for the positively selected genes, though we did observe a significant enrichment in predicted outer membrane proteins. Beyond identification of novel genes responding to host pressure, we also identified several novel sites within known AMR genes subject to positive selection. These novel sites may represent complimentary variants, emerging resistance-conferring variants or adaptive responses to the primary mutations and require further exploration.

We also considered whether variants were associated with the carriage of mobile elements or AMR through a genome-wide association analysis. We successfully identified known variants in multiple AMR-associated genes, including *gyrA* and *parC* for ciprofloxacin, *penA* for penicillin, ceftriaxone, and cefixime, *rpsJ* for tetracycline, and multiple *mtr* genes for azithromycin, validating our approach. Enticingly, we additionally identified several highly significant variants associated with AMR in genes with no known links to AMR. These include a variant in *lrgB* associated with ciprofloxacin resistance; this protein has previously been indirectly linked to ciprofloxacin resistance in *Ngo* and may play a role in the oxidative stress response^50^. Notably, we also found multiple significant variants in *mqo*, a malate-quinone oxidoreductase associated with azithromycin resistance. Despite appearing to have minimal interaction with known azithromycin targets, studies testing the efficacy of azithromycin against the protozoan parasite *Plasmodium falciparum* found reduced levels of TCA-associated metabolites, including malate, upon administration of the drug^51^. It is possible that azithromycin exerts a similar effect in gonococci, impacting its growth and that mutations in *mqo* may serve to alleviate this effect. Additional work has been conducted in *P. falciparum* to validate *mqo* as a target for drug development efforts, and our finding suggest similar efforts in *Ngo* may be pursuing^52^.

In this study we curated a representative set of 485 genetically and globally diverse gonococcal genomes from the 52,874 deposited gonococcal sequences on the NCBI SRA. Using PopNet, we assessed the population structure of our representative set, and uncovered significant relationships between population structure, geography, mobile elements, and AMR. This analysis suggests global gonococcal genetic diversity is driven by the pressure applied by antimicrobials and links the carriage of mobile elements with chromosomal variants. We used our representative set of genomes to identify putative novel phase variable genes and were able to identify phase variation of *opa* and *lgtG* potentially contributing to improved survival in the context of infection. Using a combined approach of GWAS and tests for selective pressure, we identified new sites and variants potentially associated with AMR in *Ngo*, demonstrating the utility of these large-scale analyses for studying AMR in pathogen populations.

## MATERIALS & METHODS

### Selection of sequences

An initial set of 52,874 accessions of sequencing runs were obtained from the NCBI Sequence Read Archive on September 26^th^, 2023 based on a keyword search for “*Neisseria gonorrhoeae*”. This set was reduced to 31,595 by keeping only those accessions containing paired-end Illumina sequences with associated metadata for the country and year of isolation. Additional metadata recorded when available included the site of isolation, associated BioProject, and sex of the patient. All sequence data was obtained using the SRA Toolkit v3.0.0 using the default options for the prefetch and fasterq-dump commands. Further metadata, such as minimum inhibitory concentration values were obtained from pubMLST or publications associated with the relevant BioProjects^53^. Kraken2 v2.1.3 was run to identify erroneously annotated or contaminated sequences^54^. Any sequence datasets with less than 75% of its reads annotated by Kraken as being derived from *Ngo* were discarded. Any sequences with between 75 and 85% gonococcal reads were kept if over 95% of the reads were assigned to the genus *Neisseria*.

### Variant discovery for raw sequence data

The reduced set of 31,595 isolates were aligned to the chromosomal genome of *Ngo* strain TUM19854, the current recommended reference genome for gonococci from the NCBI (obtained on October 13^th^, 2023). Alignment was performed using the Genome Analysis Toolkit (GATK) v4.2.5.0^55^. Quality control and trimming was performed using fastp v0.23.4 with minimum qualified phred quality of 20, a 40 percent limit on unqualified bases, a mean quality requirement of 20, a cut window size of 5, both front and tail cutting, and with correction enabled^56^. Trimmed reads were then passed to GATK for alignment using bwa v0.7.17 with default settings. The GATK Haplotype caller was then run with default settings to identify variants. Dependencies for GATK included samtools v1.17 and picard v2.26.3. Variants were then filtered using bcftools v1.11 and htslib v1.11 with the following parameters: allelic depth 8, allelic depth forward/reverse 3, minimum base quality 50, minimum mapping quality score 20 and percent of reads containing variant 80^57^.

### Generation of a representative set of sequences

Isolates were grouped by their country of isolation and year of isolation and the Jaccard statistic was calculated for each pair of sequences using the bedtools v2.31.0 jaccard function with default settings^58^. An initial clustering step was performed using the k-medoids fastPAM algorithm implemented in the R packages cluster v2.1.5 and factoextra v1.0.7^59–61^. For each group, a value for k was determined using the silhouette method, with the maximum value of k capped at 15. The identified medoids of the clusters were selected as representative isolates. This resulted in an initial set of 1,307 isolates. Isolates from this set were then grouped by their continent and year of isolation, and k-medoids clustering was performed as before without a cap on the value of k. Strains identified as disseminated or ocular as well as the WHO global reference collection were added to the final set, bringing the total number of isolates to 485^10,11^.

### PopNet analysis and phylogenetic tree construction

PopNet was used to cluster the reference set of genomes using the variants identified from the previous analysis^9^. A variant table was generated using the GATK VariantsToTable function from a merged vcf file of identified variants. The parameters used for the PopNet analysis were section_length= 5000, S1_iVal = 8, S1_piVal = 19, S2_iVal= 4.5, and S2_piVal= 2.5. These parameters were determined using the silhouette index readout provided by the tool. Initial phylogenetic tree reconstruction was performed with IQTree v2.2.2.7 from a sequence alignment created from the previously identified variants^62^. To begin, Model Finder Plus was run which identified TPM2u+F+I+R7 as the best substitution model^63^. IQTree was then run using this model with 1000 ultrafast bootstrap replicates^64^. All other parameters were left as default. This initial tree was used as a basis for ClonalFrameML v 1.12, which was run with the standard model with 100 pseudo-bootstrap replicates^12^. Assembled genomes were uploaded to pubMLST and automatically assigned to a cgMLST Ng_300 clade.

### Pangenome analysis

Trimmed reads were assembled *de novo* using unicycler v0.5.0 with default settings^15,65^. Gene prediction and initial annotation was performed using Prokka v1.14.5 with recommended settings and an added protein database containing annotated proteins from *Ngo* strains MS11, FA19, FA1090 obtained from the NCBI on March 7^th^, 2022^16^. The pangenome was calculated using panaroo v1.3.4 on the strict setting with otherwise default options^17^. Representative gene sequences from panaroo were supplied to eggNOG-mapper v2.19 with no restrictions on orthology identification and permitting electronic GO term annotation^66^. AMR genes were predicted from sequences uploaded to PathogenWatch on January 22^nd^, 2024^19^. The degree of resistance (resistant, intermediate, susceptible) was classified based on the EUCAST clinical breakpoints v9.0^67^. Mobile elements were predicted by BLAST+ v2.0.13 against reference sequences AY803022.1, NC_001377, NC_002098, GU479464.1, GU479465, and GU479466 which were obtained from the NCBI on December 16^th^, 2023^68^. Accessory genes were assigned to mobile elements based on a nucleotide BLAST^68^ search against this database with a minimum sequence identity of 85% and an e-value less than 1×10^-6^. The Graphia v5.0 MCL clustering tool was used to cluster accessory genes based on shared presence or absence in our representative collection with a granularity of 1.5^18^. Structures were predicted based on consensus protein sequences using AlphaFold4, and structural homology searches were performed with FoldSeek release 10^69,70^. Predicted structures were visualized with UCSF ChimeraX v1.9^71^.

### Prediction of phase variable genes

Simple sequence repeats were identified from the *de novo* assemblies and the GFF files generated by Prokka using PERF v0.4.6^20^. Genes with repeats that varied in the number of tandem repeats in more than 2 isolates where each repeat length was present in at least 5% of genomes were considered as potentially phase variable. Mutations classified as intergenic were manually verified to ensure they impacted the associated gene.

### Lower genital tract colonization for in vivo passaging

#### Bacterial growth conditions

*Ngo* MS11, parental strain, was grown on GC agar (Becton, Dickinson and Company, Sparks, USA) plates supplemented with 1% Isovitalex (ISO, Becton, Dickinson and Company, Sparks, USA) at 37°C with 5% CO_2_ for 16-20 hours. For *Ngo* isolates used for in vivo passaging, agar plates included a vancomycin, colistin, nystatin, and trimethoprim antibiotic mix (VCNT, Becton, Dickinson and Company, Sparks, USA) for *Ngo* selection.

#### Mouse strains

6-8 week old female FVB mice containing a human chromosome-derived BAC expressing CEACAM3, −5 and −6, were used for all infection experiments^25^. Animals were housed in cages with sterile rodent chow, sterile water, enrichment material, and 12-hour light-dark cycles.

#### Gonococcal *in vivo* passaging

Mice were staged by cytological vaginal smears observed under light microscopy. Those in the diestrus stage of the estrous cycle were started on an antibiotic and hormone regimen to control impact of commensal microflora on infection. Antibiotics consisted of 0.6 mg vancomycin hydrochloride (Bioshop, Burlington, ON, Canada) and 2.4 mg streptomycin sulfate (Sigma-Aldrich, Oakville, Canada) resuspended in 200 µL of PBS administered by intraperitoneal injection daily from 2 days pre-infection (−2) to 5 days post infection (+5), where an dose was provided on Day −1. Trimethoprim (Sigma-Aldrich, Oakville, Canada) was dissolved in the drinking water at 40 mg/mL for ad libitum consumption. Water soluble β-estradiol (SigmaAldrich, Oakville, Canada) was constituted in PBS (0.5 mg/200 µL), filter-sterilized and administered via subcutaneous injection on Days −2, 0 and +2.

On Day 0, bacteria were resuspended in Dulbecco’s phosphate-buffered saline (DPBS) with calcium and magnesium (PBS++, Life Technologies, Burlington, Canada)and mice were inoculated with 10^7^ bacteria in 2.5 µL Vaginal lavages were performed every two days by washing the genital tract with 10 µL of PBS++ and plated for bacterial recovery and enumeration. After ∼20 hrs bacterial isolates were frozen in 50% brain heart infusion (BHI, Becton, Dickinson and Company, Sparks, USA) and 25% glycerol (EMD Millipore, Darmstadt, Germany) and stored at −80°C. Clearance of infection was characterized by 2 consecutive vaginal lavages that tested negative for *Ngo*.

#### Gonococcal *in vitro* passaging

Five culture tubes containing 5 mL of BHI supplemented with 1% ISO was inoculated with the parental strain with a starting optical density at 550 nm (OD550) of 0.05. Bacterial cultures were placed in a 37°C shaker at 180 rpm. Every 12 hrs, bacterial stocks were generated and sub-cultured for 5 consecutive days.

#### *Ngo* whole genome sequencing

*Ngo* isolates were resuspended in 200 µL of PBS++ and genomic DNA was extracted by a QIAGEN DNeasy Blood and Tissue Kit (QIAGEN, Hilden, Germany) following manufacturer instructions. DNA quality and quantity was checked by NanoDrop ND-1000 Spectrophotometer (NanoDrop Technologies, Wilmington, USA) and Quant-iT^TM^ PicoGreen® dsDNA Assay Kit (Life Technologies, Oregon, USA) was used to quantify amount of dsDNA.Genomic DNA was prepared for whole genome sequencing by a Nextera XT library preparation kit (Illumina, San Diego, USA). Libraries were run on a 1% agarose gel to ensure libraries were of the correct length prior to poolingand submission to the Donnelly Sequencing Centre, University of Toronto for sequencing using the Illumina MiSeq platform.

### Capillary electrophoresis for confirmation of *opa* repeat lengths

Capillary electrophoresis was performed as described previously for *Neisseria meningitidis*^72^. Briefly, strains were cultured overnight on GC agar as described above, and 3 distinct colonies per plate were selected and added to 20µl of Lucigen QuickExtract buffer (Mandel, Canada). Suspended colonies were then heated at 66°C for 6 minutes, then 98°C for 2 minutes to complete the extraction. Extracted DNA was amplified with PCR using the primers in **Table 2** listed below, with the forward primer conjugated to a 6-FAM dye. Samples were then diluted to approximately 1-2ng/µl before being sent for capillary electrophoresis. The results were analyzed with the ThermoFisher MicroSatellite Analysis tool.

**Table 2:**
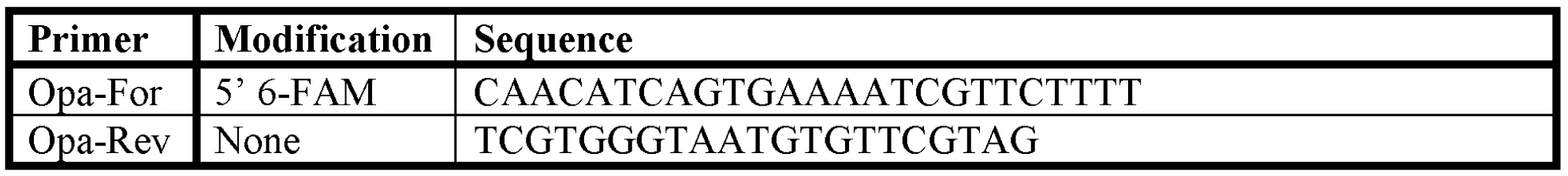
Primers used for capillary electrophoresis.

### Identification of genes under positive selection

Aligned gene sequences from panaroo for core genes with 5 or more unique sequences were used as input for selection analyses after removing identical sequences and sequences containing premature stop codons using Hyphy rmv^27^. First, a tree for each gene was built using IQTree v2.2.2.7 with 1000 ultrafast bootstraps and otherwise default settings^62^. The BUSTED algorithm was then run as implemented in HyPhy with default settings for each gene using the generated tree and panaroo alignment^28^. Genes identified as being under positive selection were then analyzed with FUBAR as implemented in HyPhy with default settings using the same inputs to identify specific sites under positive selection^29^. To assess the location of predicted residues under positive selection, structural models were generated with AlphaFold3 using the AlphaFold server based on the consensus gene sequences from panaroo^17,69^. Predicted structures were visualized with UCSF ChimeraX v1.9^71^.

### Genome wide association study

The genome wide association study was performed using pyseer with the linear mixed model setting using the VCF file generated as described above^39^. The distance matrix was calculated using the ClonalFrameML generated tree described above. All other pyseer settings were kept at default. The count_patterns script was used to establish a significance threshold for each tested phenotype. The impact of each variant was predicted with SnpEff using the *Ngo* TUM19854 reference genome obtained as previously described^73^. Structural alignments and comparisons were performed using TM-align with default settings^40^.

## ACKNOWLEDGEMENTS

We are grateful to Odile Harrison, University of Oxford, for helpful advice and technical support, and the Division of Comparative Medicine & Donnelly Sequencing Centre at the University of Toronto for technical support. This study was supported by operating grants from the Canadian Institutes for Health Research (PJT-159677) and National Institutes of Health (1R01AI146941-01A1). S.D.G. is supported by the Canada Research Chairs Program. D.C. is a recipient of the EPIC Doctoral Award and the DSI Doctoral Fellowship.

## ETHICS STATEMENT

All animal procedures conducted in this study were approved by the Local Animal Care Committee (LACC) at the University of Toronto (Protocol #20011775), which is in compliance with ethical and legal requirements under Ontario’s Animals for Research Act and the federal Canadian Council on Animal Care.

## DATA & CODE AVAILABILITY

The data sets generated in this study are provided in the Supplementary Information and Source Data files. Codes for analysis and data are available on Github [https://github.com/ParkinsonLab/Neisseria_work.git].

## AUTHOR CONTRIBUTIONS

J.P. and S.D.G conceived the idea of the research and supervised it. D.C and E.J. performed the computational and data analysis. D.C. and J.M. conducted the mouse passaging and phase variation experiments and analyzed the data. D.C. wrote the manuscript with contributions from J.P., S.G.D. & J.L.

## SUPPLEMENTARY INFORMATION

**Supplementary Figure S1: Overlap of clustering from PopNet, cgMLST & PopPunk.** Tree (left) from ClonalFrameMLST overlaid with PopNet clusters, 11 largest clusters from cgMLST and PopPunk. Network view (right) generated by PopNet overlaid with 11 largest clusters from cgMLST and PopPunk.

**Supplementary Figure S2: Predicted AMR from PathogenWatch.** Predictions based on assembled genomes using PathogenWatch. Degree of resistance based on EUCAST clinical breakpoints v9.0.

**Supplementary Figure S3: Colonization of female transgenic FVB mice by *in vivo* passaged *Ngo*.** Mice were checked every 2 days after 1-day post-infection for evidence of colonization. Mice contained a human chromosome-derived BAC expressing CEACAM3, −5 and −6.

**Supplementary Figure S4: Variants associated with AMR identified through GWAS.** Manhattan plots of variants with significant associations with penicillin (A), tetracycline (B), ceftriaxone (Ci), and cefixime (Cii) resistance. Only variants with an LRT p-value <0.05 are shown. Red lines indicate significance thresholds determined by Bonferroni correction with the number of unique variant patterns as the number of multiple tests.

**Supplementary Table S1: SRA metadata from initial set of 31,595 gonococcal genomes.**

**Supplementary Table S2: Isolate metadata for representative set of 485 genomes.**

**Supplementary Table S3: Metadata for all genes identified within the gonococcal pangenome.**

**Supplementary Table S4: Clusters of accessory genes determined through Graphia MCL.**

**Supplementary Table S5: Phase variable genes identified through genetic screening of representative genomes.**

**Supplementary Table S6: Evidence for signatures of positive selection at the gene and residue level.**

**Supplementary Table S7: Variants significantly associated with AMR identified through GWAS.**

