## Supplementary figures and images for "Network-based population analysis of 31,595 gonococcal genomes reveals phase variability, genetic diversity and mobile element dynamics drive antimicrobial resistance and phenotypic diversity"

### Supplementary Figure S1

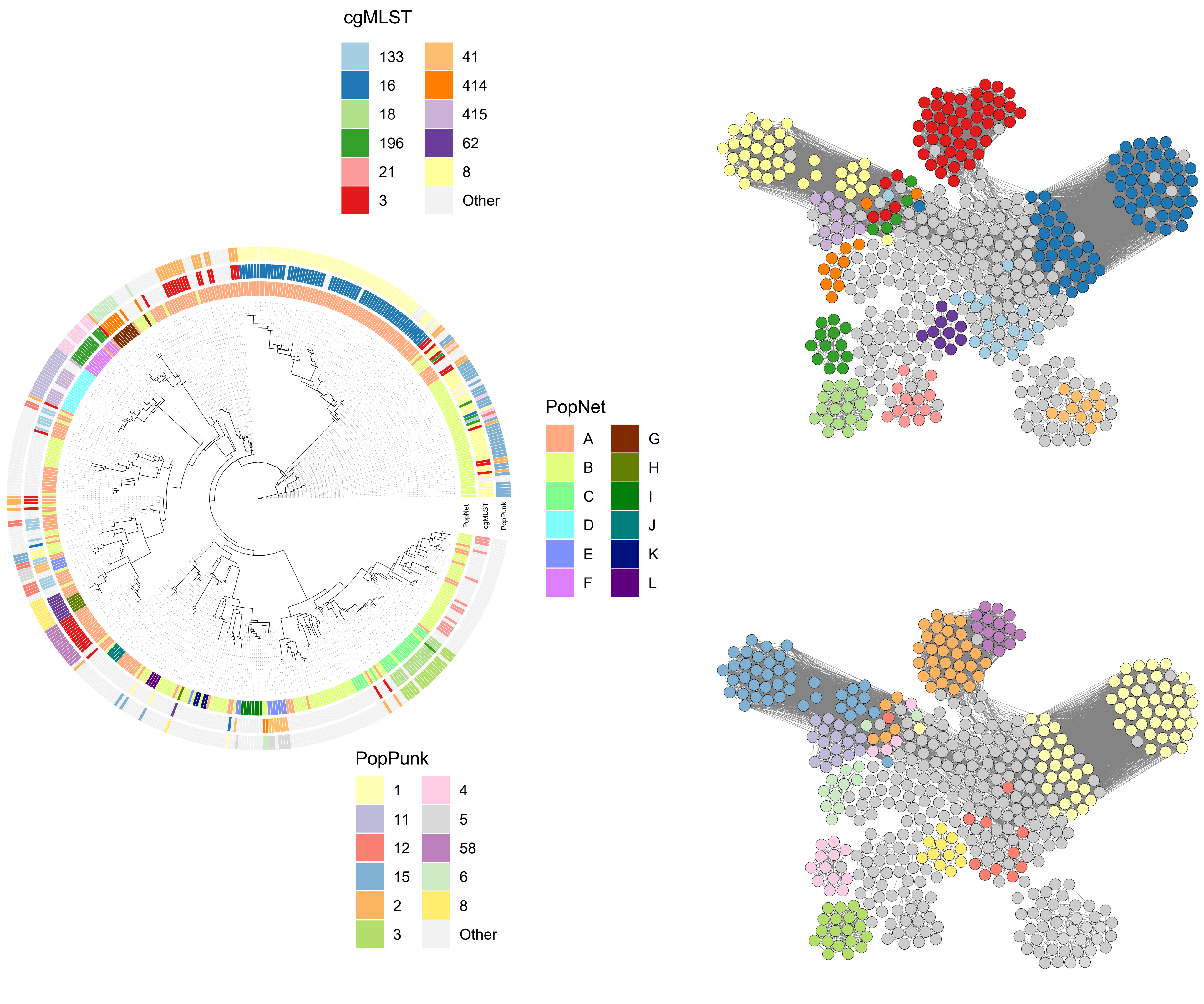

### Supplementary Figure S2

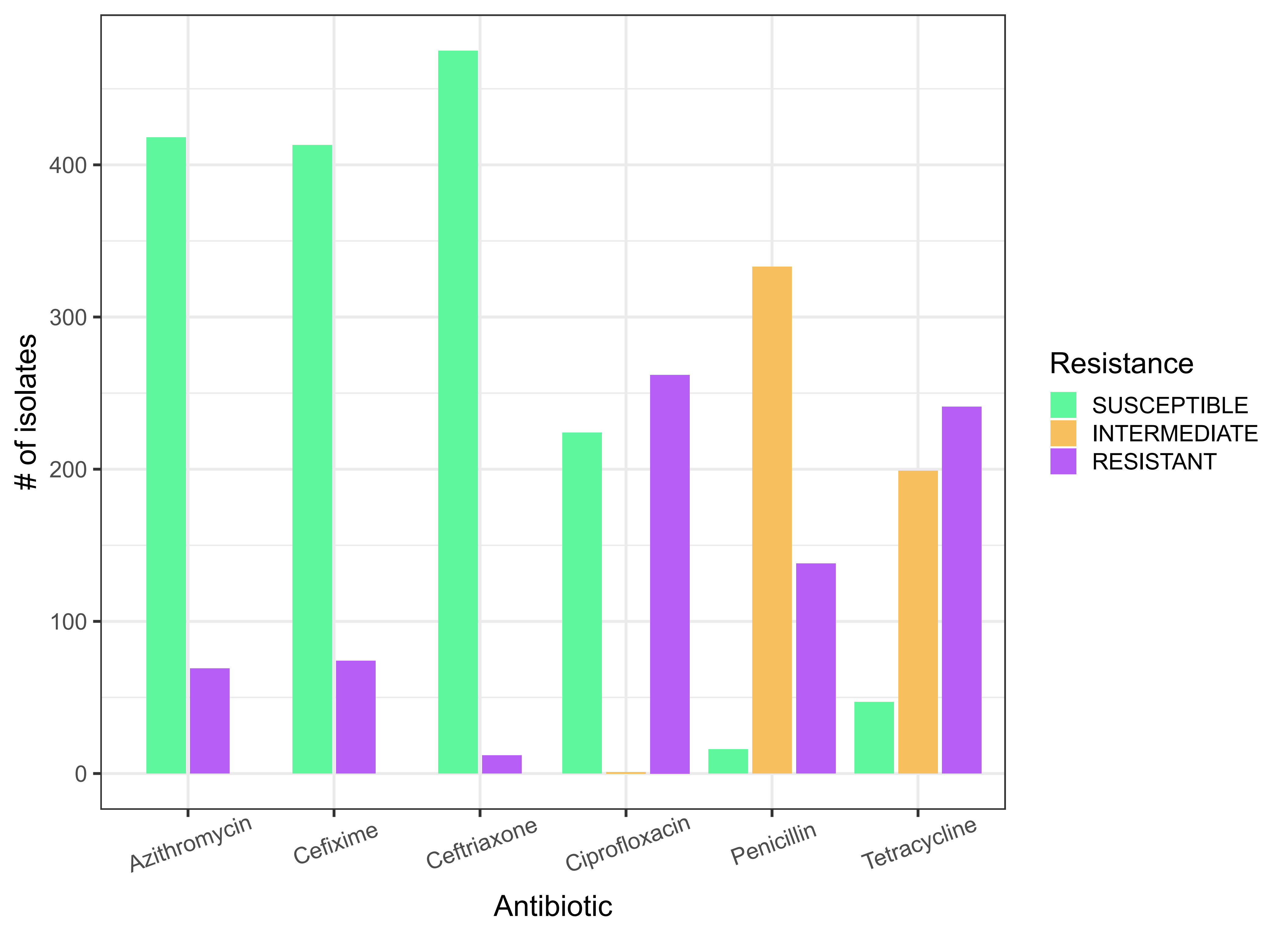

### Supplementary Figure S3

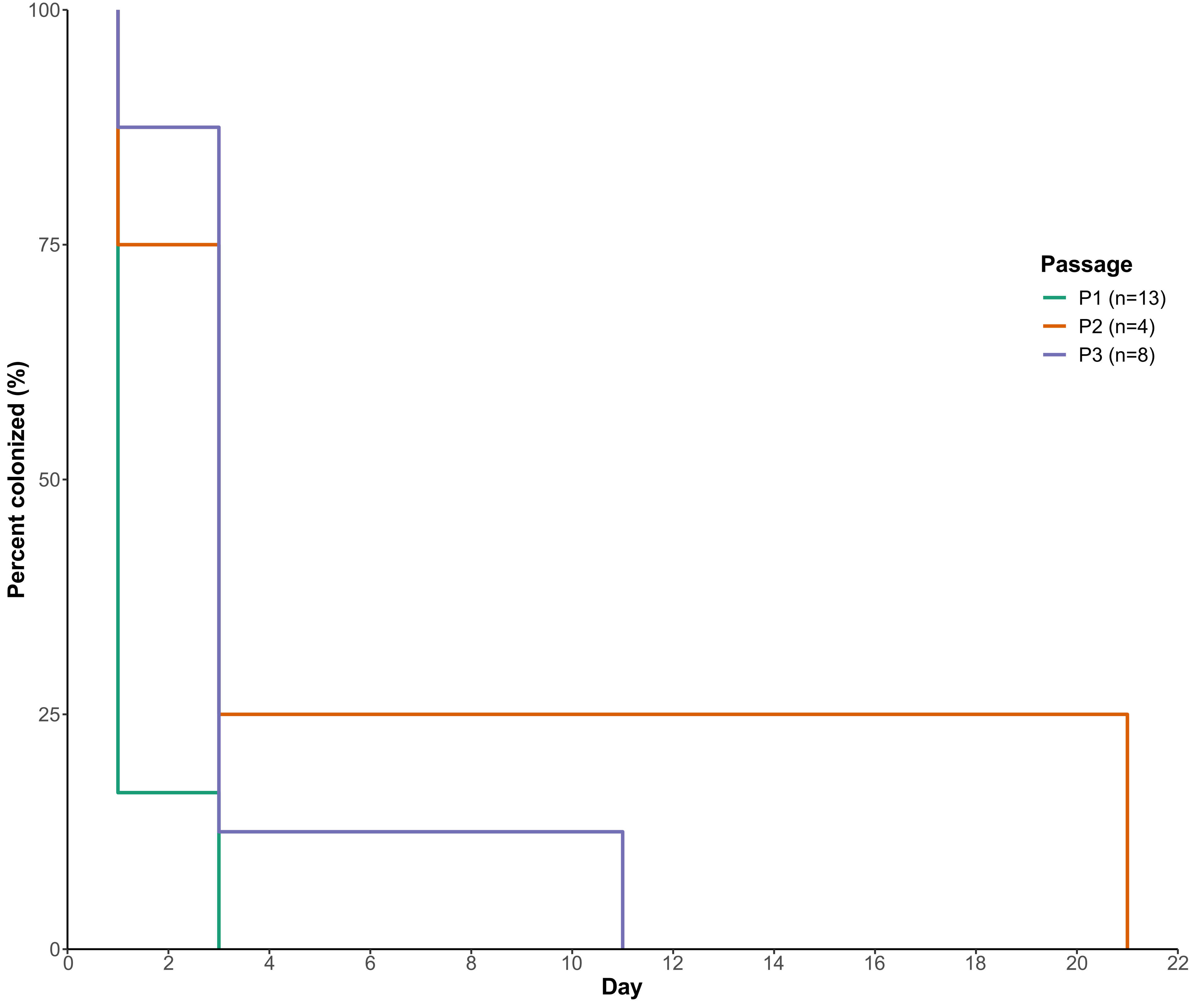

### Supplementary Figure S4

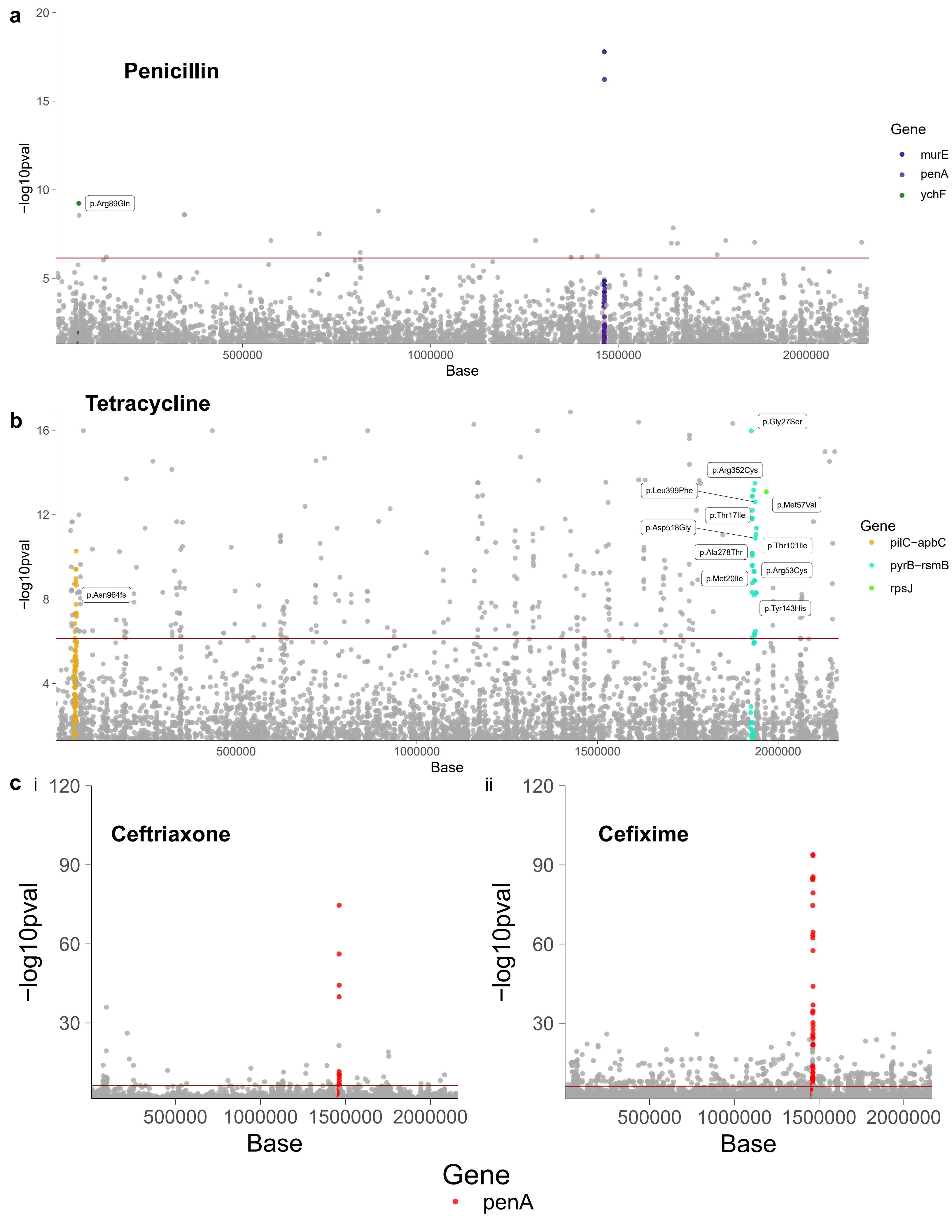
